# A Field-Based Study of Phyllosphere Mycobiomes in Apple Orchards Under Varying Agricultural Management Strategies

**DOI:** 10.1101/2025.09.25.678622

**Authors:** Sophie Boutin, Jonathan Rondeau-Leclaire, Alexis Roy, Isabelle Laforest-Lapointe

**Affiliations:** Département de biologie, Université de Sherbrooke, Sherbrooke, QC J1K 2R1, Canada; Centre SÈVE, Université de Sherbrooke, Sherbrooke, QC J1K 2R1, Canada

**Author notes:** Co-first authors (Sophie Boutin, Jonathan Rondeau-Leclaire). Corresponding author: Isabelle Laforest-Lapointe.

**Keywords:** mycobiome, apple tree, *Malus domestica*, phyllosphere, agricultural management strategies, microbial ecology, ITS amplicon sequencing, on-farm study

## Abstract

Microbial communities in the phyllosphere are key players in plant health and disease resistance, yet their response to agricultural management strategies remains poorly understood under field conditions. Here, we compare fungal community composition and diversity across conventional and organic apple orchards using ITS amplicon sequencing. Leaf samples were collected from six sites at three distinct time points during the 2023 growing season (in May, July, and August) corresponding approximately to monthly intervals throughout the summer. Flower samples were collected from the same trees in May. Our analyses reveal that agricultural management strategies are significantly associated with fungal community structure, with effects intensifying from May to July. Both types of management strategies showed enrichment for different genera known to include common apple tree pathogens: *Alternaria* and *Podosphaera* were associated with conventional sites, while *Didymella* and *Ramularia* were associated with organic sites. Although fungal alpha diversity was higher in May at conventional orchards compared to organic orchards, it declined over time at conventional sites while it remained stable at organic sites. Together, these patterns indicate that distinct management interventions impose contrasting selective pressures on the apple tree phyllosphere mycobiome, thus shaping both broad fungal community composition and the dominance dynamics of key fungal taxa. Our findings underscore the ecological relevance and inherent challenges of field-based microbiome research, and provide insights to inform the development of sustainable orchard management strategies grounded in fungal community dynamics.

## Introduction

As the world population keeps growing, agriculture faces increasing pressure not only to meet the rising food demand, but also to ensure efficient, stable, and sustainable production systems that can support long-term productivity under variable environmental conditions. In parallel, pressures on ecosystems biodiversity and functions have never been so strong, and global change is deeply impacting biomes’ temperature and precipitation regimes [1]. Now more than ever, crop producers face the heightened challenges of optimizing land use and limiting the spread of plant bacterial and fungal diseases, while navigating social expectations to minimize local environment disruptions. However, crop management often relies on the use of chemicals such as antibiotics, fungicides, and insecticides to maintain productivity. These products can be damageable to the environment, potentially leading to water pollution [2] and to the emergence of resistances in microbial pathogens [3, 4].

While the use of chemicals in agriculture provided crucial benefits to human society in terms of food security during the Green Revolution of the 1950s, contemporary research is now suggesting that many pesticides, whether of synthetic or natural origin, disrupt the overall plant-associated microbial communities [5, 6]. These disturbances in soil and plant microbiomes, even at low doses, may ultimately affect plant fitness and broader ecosystem functions [7]. As a result, alternative management strategies (e.g., organic cultures) are being promoted for their potential to reduce synthetic chemical inputs as well as preserve ecosystem biodiversity and functions [8] even if they can be less productive per land area in the short term. Yet, organic crop management is not free from chemical disturbances. The recurring application of biological and mineral-based plant protection products (e.g., copper, sulfur, microbial biocontrol products such as the yeasts *Aureobasidium* spp. contained in “Blossom Protect” [9] and the bacteria *Bacillus thuringiensis* contained in “Bt” [10]) can also impact microbial communities associated with crops.

Plant flowers, leaves, and roots harbour diverse microbial communities (i.e., assemblies of microorganisms including bacteria and fungi) that can impact plant host productivity and general health [11, 12]. For instance, members of the *Arabidopsis* microbiota can antagonize the widely distributed foliar pathogen *Pseudomonas syringae* [13], and plant growth promoting rhizobacteria (PGPR) are foreseen to be key players in plant growth [14, 15]. Rhizosphere (i.e., root-associated) microbial communities are generally well-characterized and are known to respond strongly to soil properties, root exudates, and agricultural management strategies [16, 17]. For example, perennial crops such as grapevine and olive show consistent enrichment of PGPR and mycorrhizal fungi under reduced chemical inputs [18, 19]. In contrast, the phyllosphere (i.e., flowers and leaves) microbial communities appear to be mostly influenced by aerial and vector-based (e.g., insects) dispersal, leaf surface chemistry, and microclimatic conditions [11], leading to higher stochasticity and sensitivity to canopy-level management [20]. These broad ecological differences suggest that patterns observed in the rhizosphere, such as strong nutrient-driven selection, may not apply to the phyllosphere, where dispersal limitation and environmental filtering more likely dominate. In the last decades, the phyllosphere has gained scientific interest due to its demonstrated impacts on general plant health [11].

Fungi, although expected to be less abundant than bacteria in the phyllosphere [11], are known to affect host plant and ecosystems through drought tolerance improvement [21], pathogen regulation [9], and leaf litter decomposition [22]. Several pathogenic fungal taxa are present in the apple tree phyllosphere including *Venturia inaequalis* (the causal agent of apple scab) [23], *Podosphaera leucotricha* (the causal agent of powdery mildew) [24], and *Alternaria* spp. (the causal agent of *Alternaria* leaf blotch) [25]. For example, *V. inaequalis* is a well-documented and economically significant pathogen in apple orchards worldwide, and its presence is expected in apple tree phyllosphere communities, particularly under conditions favorable to disease development such as cool and moist spring weather [23]. Yet, the apple tree phyllosphere is also colonized by a broad diversity of commensal and beneficial fungal taxa [26]. In a previous study, our group identified *Aureobasidium*, *Sporobolomyces*, and *Mycosphaerella* as the three main fungal genera observed in the apple tree phyllosphere [26]. Our work also showed that site and time are strong drivers of microbial (both bacteria and fungi) community composition and diversity in field conditions [26, 27]. Yet, in Boutin *et al.* (2024) [26], when stratifying our data by agricultural management strategy, the organic site showed no significant difference in alpha diversity across the three time points, in sharp contrast with the observable trend at the two conventional sites. Therefore, to broaden the impact of our research and investigate this discrepancy, we extended our sampling design of the apple tree phyllosphere to include more sites of varying agricultural management strategies.

While organic and conventional systems differ in their management strategies, it is important to note that most economically viable agricultural systems still rely on plant protection measures including fungicidal and insecticidal compounds [28]. The primary distinction lies in the active ingredients’ origin (natural vs. synthetic) and the regulatory frameworks governing their use [29]. In organic systems, management strategies include the use of broad-spectrum fungicides and bactericides such as copper, which has a well-documented ecotoxicological profile, including persistence and toxicity to non-target organisms. Yet, the continued use of copper reflects its long-standing utility rather than environmental benignity [30]. These contact fungicides remain susceptible to wash-off and therefore exhibit limited rainfastness, often necessitating repeated applications due to non-systemic mode of action and surface localization [31]. Because these treatments act broadly on exposed microbial communities (non-targeted effects), they may also disturb repeatedly plant leaf microbial communities, thus contributing to non-specific reductions in epiphytic microbial diversity [32]. In contrast, conventional management strategies largely rely on synthetic plant protection products (often with systemic or partially systemic effects) which have more specific modes of action, the capacity to persist within plant tissues, and act as strong ecological filters [28]. This internal activity allows for longer intervals between applications because of the products’ prolonged efficacy, its protection of newly developed tissues, and its reduced susceptibility to wash-off. Therefore, these treatments could lead to fewer repeated disturbance events of plant-associated microbial communities compared to surface-acting contact fungicides [30, 33]. Thus, rather than assuming that organic management is intrinsically more compatible with leaf microbiome integrity, it is essential to evaluate how different management strategies shape plant-associated microbial communities. In this context, research on the effects of agricultural management strategy (conventional and organic) on plant-microbe interactions in field is crucial to advance our knowledge of these ecosystems and better support crop producers.

The current study focuses on the apple tree (*Malus domestica*, Borkh.) in commercial orchards, to explore how canopy-level management, site identity, and time of sampling shape phyllosphere fungal communities. To disentangle the relative importance of agricultural management from other sources of variability, it is key to consider fungal community dynamics across spatial and temporal gradients under operational conditions. In this work, we leveraged a multi-site and multi-temporal sampling design previously established [26, 27] to assess how fungal communities associated with apple tree leaves and flowers differ between contrasting management strategies (conventional vs. organic), while accounting for variation among orchards and across the growing season. This approach allowed us to evaluate whether management-associated differences are consistent across sites or contingent on local conditions and broad phenological timing. In addition, because genotype has been shown to influence the common apple tree leaf microbial assembly processes [34] and resistance to foliar pathogens [35], we considered the potential contribution of apple cultivar identity to phyllosphere community structure. Although cultivar effects are generally expected to be weaker than those of management, site, or season, incorporating this factor enables a more comprehensive understanding of the drivers shaping the apple phyllosphere mycobiome.

Here, our main objectives were to (1) assess the associations between agricultural management and fungal community composition, (2) characterize how alpha diversity varies across time, sites, and agricultural management strategy, and (3) identify differentially abundant taxa between conventional and organic sites. These analyses were performed on a large dataset including four to seven cultivars at each site, as well as a smaller subset of two cultivars present at every sites. Building on previous results from our group [26, 27] and others (**Table S1**), we hypothesized that fungal community composition would diverge between agricultural management strategies, with these differences reflected in the differential abundance of specific taxa. We further expected that the use of synthetic chemicals at conventional sites would impose stronger selective pressures, leading to greater shifts in the composition of dominant fungal taxa, including key apple leaf pathogens such as *V. inaequalis*. In addition, we predicted that leaf fungal alpha diversity (Shannon index) would be lower at conventional sites. Finally, we anticipated significant spatial (site-specific) and temporal variation in fungal community composition and diversity, driven by taxon-specific responses to climatic conditions and management strategies, as well as dispersal from local reservoirs (e.g., soil, surrounding vegetation, and land use).

## Materials and methods

### Sampling

Six commercial orchards (A-B1-B2-C-D1-D2) from the Eastern Townships region of Québec (Canada) were sampled in 2023. These orchards (sites) were selected based on available cultivars, agricultural management strategy (conventional or organic), and spatial proximity. All organic orchards included in this study complied with the Canadian *Ecocert* certification standards for organic production. This certification prohibits the use of synthetic fertilizers, synthetic pesticides, and genetically modified organisms, while also restricting crop protection and soil amendments to a list of approved substances [36]. Two pairs of orchards (B1-B2, D1-D2) were selected as pairs of contrasting management strategy with spatial proximity. Sites B1 and B2 are respectively conventional and organic orchards situated less than a kilometer apart. Sites D1 and D2 are conventional and organic plots within the same commercial orchard. These two pairs of sites allowed us to test for differences in agricultural management strategy while controlling for the effect of distance. For climate, the mean daily temperatures at the closest meteorological station ranged between 1.7°C and 24.3°C from May 1^st^ to August 31^st^. On May 17^th^, there was an extreme frost event in the region. The total precipitation in May, July, and August was respectively of 45, 310, and 140 mm. More details about the sites, the sampled cultivars, the mean daily temperatures, and total precipitations are available in **Table S2** and **Figure S1**.

At each site, three trees from each cultivar were randomly selected. In May, during the flowering period (stage 6) [37], leaf and flower samples were collected. Producers were asked to notify us when flowers reached BBCH [38] stage 65 (full bloom), while ensuring compliance with safety regulations governing human access to orchards during and after pesticide applications. In July and August, which correspond to stage 7 and 8 respectively [37], only leaf samples were collected as flowering season was over. On the same day as leaf and flower collection, samples were stored at -20°C in the laboratory until cell harvesting and DNA extraction.

### Fungal cells collection and DNA extraction

Using the method previously described by Boutin *et al.* [26, 27], samples were thawed at room temperature approximatively one hour before proceeding to cell collection. Then, 100 ml of 1X Redford Buffer [39] was added to each sampling bag. The bags were shaken during five minutes, then the buffer was decanted equally into two 50 ml tubes. The wash-off approach was selected to characterize the broader community of leaf-associated fungi present on the phyllosphere, which includes epiphytes as well as taxa capable of transitioning to or from an endophytic lifestyle. This approach may therefore overrepresent epiphytic taxa, including surface-growing pathogens.

The tubes were then centrifuged at 4000 g for 20 minutes and the supernatants were discarded. The pellet of the first tube was resuspended in 800 µl of solution CD1 from the DNeasy^®^ PowerSoil^®^ Pro Kit (QIAGEN). The content of this tube was transferred to the second tube to resuspend its pellet. Up to 1 ml was transferred to a bead beating tube, which was kept at -20°C until DNA extraction. Before DNA extraction, bead beating tubes containing the microbial pellets and CD1 were thawed at room temperature. Tubes were placed in a Bead Mill Homogenizer (Fisher) set to the following parameters (speed: 3.25 m/s; time: 2 minutes; cycles: 5; breaks: 30 seconds). The rest of the extraction followed the quick start protocol and DNA was eluted in 75 µl of solution C6. Extracted DNA was stored at -20°C until it was sent to the sequencing platform.

### Sequencing

The ITS1 region was amplified using the ITS1F (5’-CTTGGTCATTTAGAGGAAGTAA-3’) [40] and ITS2 (5’-GCTGCGTTCTTCATCGATGC-3’) [41] primers. Samples were sequenced at the CERMO genomic platform (Université du Québec à Montréal) on the Illumina MiSeq platform at 2×300 bp.

### Amplicon sequence variants processing

A total of 9,333,318 sequences from 364 samples were processed using the DADA2 pipeline v1.34 [42] in R v4.4.0 [43] using default parameters unless otherwise stated. Briefly, low complexity sequences and those containing ambiguous nucleotides were discarded using the *filterAndTrim* function with *rm.lowcomplex = TRUE* and *maxN = 0*. Primers were removed using cutadapt v2.10 [44] with parameters *-n 2 -m 21 -M 300*, then sequences were quality-filtered using *filterAndTrim*, allowing a maximum expected error *maxEE = c(4, 4)* and a minimum length *minLen = 100*. After learning the error model, sample compositions were inferred using *learnErrors(pool = “pseudo”).* Forward and reverse sequences were merged, then bimeras removed using *method = “consensus”*. The taxonomy of the resulting amplicon sequence variants (ASVs) was assigned using the UNITE ITS database v10.0, 04-04-2024 [45]. ASVs with fewer than a total of 10 sequences across all samples or with no taxonomic classification at the Phylum level were removed. Then, samples with fewer than a total of 2,000 sequences were discarded, resulting in a total of 329 samples kept for analysis (the 35 discarded samples are listed in **Table S3)**. This resulted in a total of 5,022,964 total sequences (mean 15,267 ± 8,189 per sample) and 2,812 ASVs (mean 216 ± 144 per sample). On average, each ASV was found in 25 ± 39 samples. Sample metadata, sequence and taxonomy tables were built into a phyloseq object (phyloseq v1.46.0 [46]) for further analyses. For alpha and beta diversity analyses (see below), samples were rarefied to 2,500 sequences to control for uneven sequencing depth using the *rarefy_even_depth* function from the vegan package v2.7-1 [47]. This resulted in a total 822,500 sequences and 2,795 ASVs (mean 146 ± 112 per sample). On average, each ASV was found in 17 ± 31 samples after rarefaction (**Table S4**).

### Statistical analyses

Statistical analyses were performed using base R functions and the rstatix v0.7.2 and vegan packages. Plots were generated using the ggplot2 v3.5.2, ggpubr v0.6.1, ggrain v0.0.4, ggh4x v0.3.1, see v0.11.0, and patchwork v1.3.1 packages.

For each dataset, we analyzed alpha diversity based on the Shannon index and beta diversity based on the Bray-Curtis dissimilarity. Shannon diversity indices were calculated using the *estimate_richness* function from phyloseq. Beta diversity was estimated by first variance-stabilizing the rarefied count tables using the *varianceStabilizingTransformation* function from DESeq2 v1.48.1, then computing the Bray-Curtis dissimilarity index using the *vegdist* function from vegan. Permutational multivariate analyses of variance (PERMANOVA) were applied to the resulting dissimilarity matrices using the *adonis2* function from vegan with 999 permutations. Homogeneity of group dispersions was verified using the *betadisper* function (**Table S5**). The dissimilarity matrices were visualized using principal coordinates analysis (PCoA). Aggregate sample composition bar charts are based on the mean relative abundance of each taxon, showing the top 20 taxa that were most relatively abundant on average.

### Flower fungal beta and alpha diversity

To characterize the fungal communities associated with apple tree flowers, we analyzed a subset of 54 flower samples (35 from conventional and 19 from organic orchards), collected across four sites (A, B1, B2, and C). Sites D1 and D2 were excluded due to the absence of flowers at the time of sampling.

Beta diversity dynamics were tested using the following model: [*Community ∼ (Management/Site) × Cultivar*] with ‘Site’ nested in ‘Management’ to reflect the sampling design. A Shapiro-Wilk test revealed that alpha diversity was not normally distributed; therefore, the Kruskal-Wallis non-parametric test was used to infer whether organic and conventional sample diversity originated from the same distribution.

### Leaf fungal beta diversity

To assess the influence of agricultural management strategy on the fungal communities associated with apple tree leaves, we analyzed all 275 leaf samples, encompassing all sites and cultivars. Beta diversity dynamics were tested using the following model: [*Community ∼ (Management/Site) × Time + (Management/Site) × Cultivar*] with ‘Site’ nested in ‘Management’ to reflect the sampling design. Permutations were constrained within individual trees (TreeID) to account for repeated measures. Separate analyses were also conducted for the Honeycrisp (n = 51) and Spartan (n = 53) cultivars to confirm that our results were robust despite uneven cultivar representation across sites. To control for temporal effects, we repeated the PERMANOVA and PCoA independently for each sampling time point (May, July, August), using the model: [*Community ∼ (Management/Site) × Cultivar*], allowing to estimate the variance explained by agricultural management strategy at each time point. To further highlight site-specific patterns, ordinations were visualized to highlight pairs of sites that were spatially closed to one another: (1) for sites B1 and B2 as well as (2) for sites D1 and D2. To visualize the variation in fungal class composition across the PCoA ordination, we performed an *envfit* analysis from the vegan package to test for correlations between the relative abundance of fungal classes and the ordination axes. Significance was assessed using permutation tests, and a Bonferroni correction was applied to account for multiple comparisons.

### Leaf fungal alpha diversity

Because a global linear model across time points violated key assumptions (non-normality, heteroscedasticity, and temporal autocorrelation; Shapiro-Wilk, Breusch-Pagan, and Breusch Godfrey tests), leaf fungal alpha diversity was analyzed using Kruskal-Wallis non-parametric tests, followed by post hoc pairwise Dunn’s tests with Bonferroni correction. These analyses were performed to assess variation in diversity across sampling time points (May, July, August) within each site, across time points within each agricultural management strategy, and across strategies at each time point.

All analyses (for composition and alpha diversity) were also performed separately for the Honeycrisp (n_Flowers_ = 8; n_Leaves_ = 51) and Spartan (n_Flowers_ = 9; n_Leaves_ = 53) cultivars to confirm that our results were robust to uneven cultivar representation across sites.

### Differential absolute abundance analyses

Differences in absolute abundance between organic and conventional sites were estimated using the log-linear model implemented in ANCOM-BC2 [48] and are expressed as natural log fold differences (LFD) in absolute abundances. The following procedure was done separately for flower and leaf samples. First, non-rarefied sequence counts were aggregated at the genus level, discarding any ASV with no taxonomic assignment at that level. Genera found fewer than 10% of samples (6 flower and 28 leaf) were not considered. The model controlled for cultivar, as well as sampling month in the case of the leaf samples. *P*-values were manually corrected using the Holm procedure, because ANCOM-BC2’s *p*-value correction only accounts for the number of tests done in a single run, while the command was executed once for leaf and once for flower samples. Genera with an adjusted p-value < 0.01 and having passed the pseudo-count sensitivity check were considered as differentially abundant (enriched) in either group. Only genera with an absolute LFD > 1 were reported.

## Results

### Flower samples across conventional vs. organic sites

We first explored the impact of agricultural management (conventional vs. organic) on flower fungal community composition and diversity. From the principal coordinates analysis (PCoA), we observed that samples from conventional and organic management strategy clustered separately (**Figure 1A**). The permutational variance analysis (PERMANOVA) revealed that agricultural management strategy explained up to R^2^ = 20% (*P* = 0.001) of the variation observed within those samples (**Table 1**). Site identity was also a driver of fungal community structure, explaining R^2^ = 13% of the variation (**Table 1**). In terms of alpha diversity, flower samples from organic sites had a lower mean Shannon index than those from conventional sites (conventional = 4.3, organic = 3.7; *P* < 0.0001) (**Figure 1B**). The three main fungal genera found in samples from conventional sites were *Cladosporium*, *Aureobasidium*, and *Fomes* which respectively represented 8%, 8%, and 7%, summing up to 23% of sequences. At organic sites, the main fungal genera were *Ramularia, Didymella, Gyoerffyella*, representing 17%, 17%, and 9%, summing up to 43% of sequences (**Figure 1C**).

**Figure 1.**
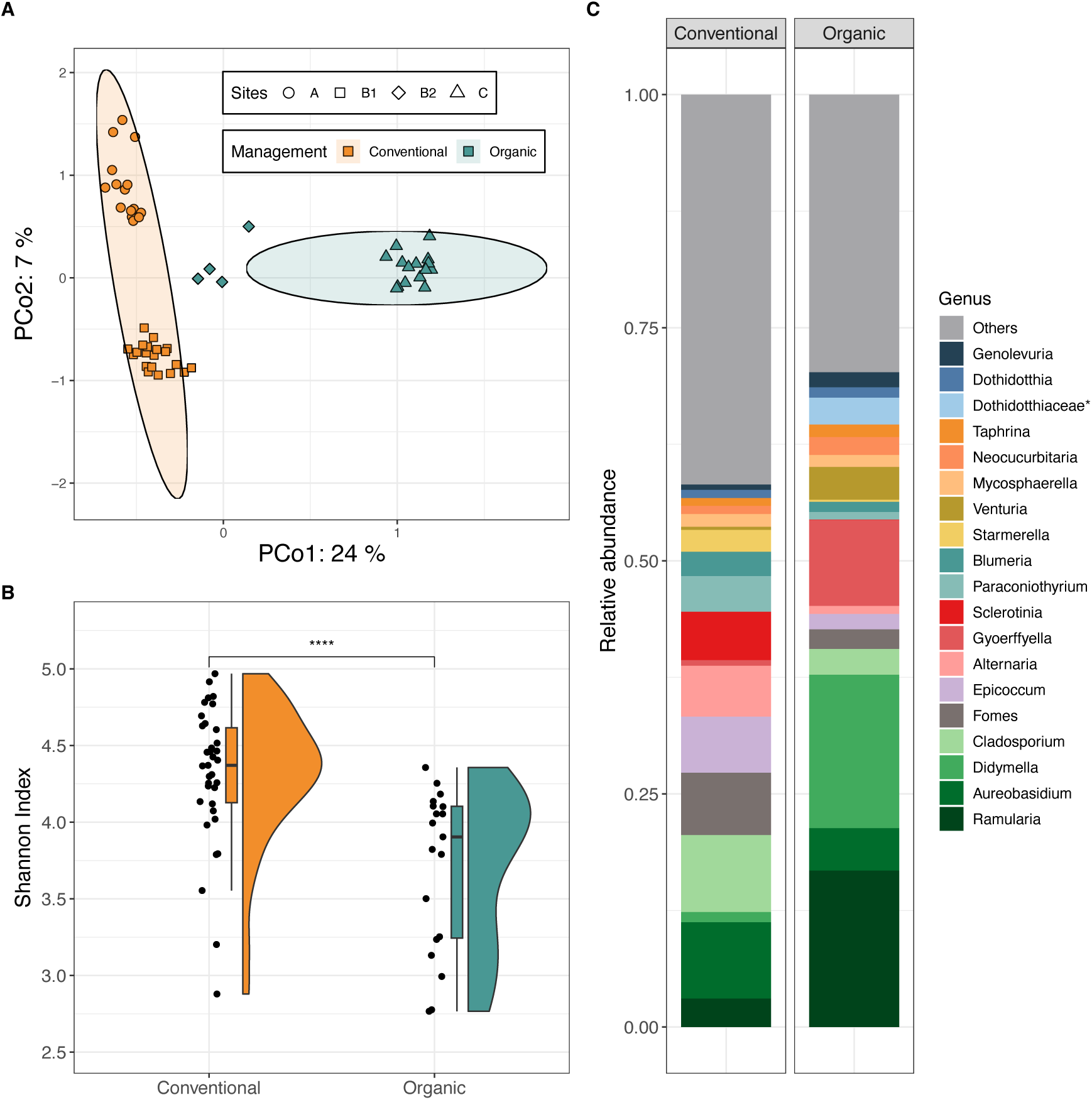
Fungal composition and diversity of flower samples. **(A)** Principal Coordinates Analysis (PCoA) based on Bray-Curtis dissimilarities among flower samples. Point shapes and colors indicate sampling sites and agricultural management strategies. Ellipses represent 95% confidence intervals. **(B)** Shannon alpha diversity index across conventional (n =35) and organic (n = 19) sites. Asterisks indicate significance levels from Wilcoxon-Mann-Whitney test (**** P ≤ 0.0001). **(C)** Average relative abundance of the 20 most abundant fungal genera across flower samples. Taxa not among the top 20 are grouped under “Others.” Bar charts are separated by agricultural management strategy. *Asterisks denote taxa identified at the family level but with unknown genus*.

**Table 1.** Factors explaining the variation within all flower samples (PERMANOVA on Bray-Curtis dissimilarities)

|  | R<sup>2</sup> (%) | F | Pr(>F) |  |
| --- | --- | --- | --- | --- |
| Management strategy | 20.2 | 16.9 | 0.001 | *** |
| Cultivar | 12.4 | 1.5 | 0.008 | ** |
| Management:Site<sup>*1</sup> | 12.7 | 5.3 | 0.001 | *** |
| Management:Cultivar | 8.9 | 1.2 | 0.058 | ns |
| Management:Site:Cultivar | 7.7 | 1.3 | 0.050 | * |
| Residuals | 38.1 |  |  |  |
| Total explained | 61.9 |  |  |  |

<sup>\*1</sup> Site is nested within Management.

We then sought to identify genera enriched in samples from either organic or conventional sites. To that end, we estimated fold differences in absolute abundances for each genus using a log-linear model (ANCOM-BC2), controlling for cultivar type (**Figure 2**). In flower samples, eight and twenty-six genera were found to be differentially enriched (adjusted p < 0.01) in conventional and organic sites, respectively. Lecanoromycetes (n = 2 genera) and Eurotiomycetes (n = 1 genus) were only enriched in conventional sites, while Agaricomycetes (n = 11 genera), Phaeococcomyces and Cystobasidiomycetes (n = 1 genus each) were only enriched at organic sites. The most differentially enriched genus, *Exidia*, was estimated to be almost 37 times more abundant (3.6 ± 0.2 LFD) in organic sites compared to conventional sites. Of note, the genera *Podosphaera* (e.g., *P. leucotricha*, the causal agent of powdery mildew) and *Penicillium* (e.g., *P. expansum*, causing blue mold rot) were more abundant on flowers at conventional sites (2.7 ± 0.2 LFD; **Figure 2**). In contrast, the genera *Venturia* and *Alternaria* were not differentially abundant across management strategy although they showed higher relative abundance on average respectively at organic and conventional sites (**Figure 1**).

**Figure 2.**
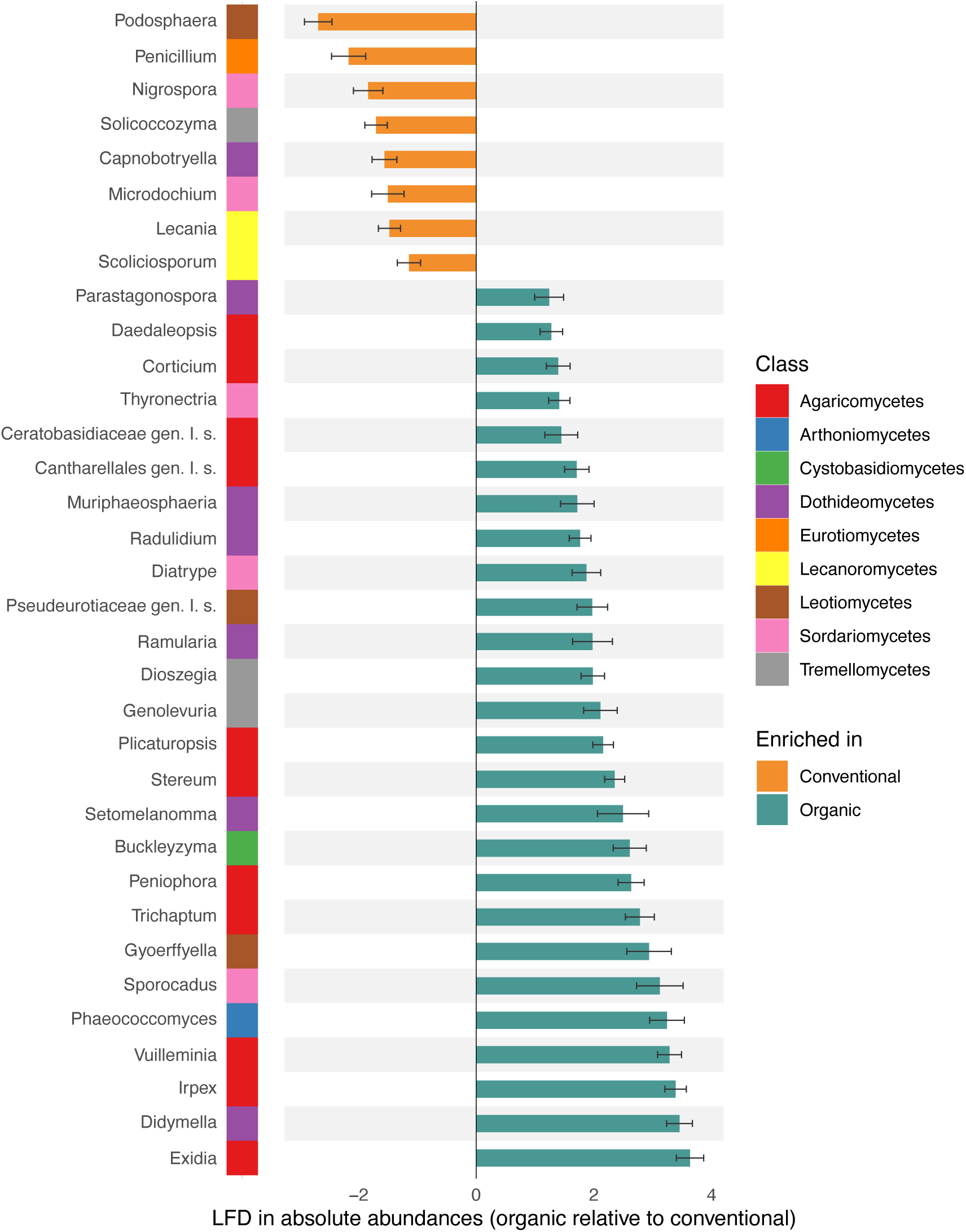
Differential abundance analysis of flower communities comparing differences in absolute abundance of genera across agricultural management, controlling for cultivar. A negative natural log fold difference (LFD) corresponds to an enrichment in the conventional sites (i.e., the reference group in the log-linear model), whereas a positive LFD corresponds to an enrichment in the organic group. Gen. I. s.: *Genus Incertae sedis*.

Overall, we obtained similar results when looking at flowers from Honeycrisp and Spartan cultivars only. Indeed, samples clustered separately by management strategy (**Figure S2A**). Of note, management strategy explained an even greater amount of the variation observed (R^2^ = 29%, *P* = 0.001) within samples (**Table S6**), and the difference in alpha diversity became marginally statistically significant (conventional = 4.4, organic = 3.9; *P* = 0.07; **Figure S2B**). The two most relatively abundant genera (*Ramularia* and *Aureobasidium*) (**Figure S2C**) were the same as for the whole dataset.

### Leaf samples across conventional vs. organic sites

The impact of agricultural management strategy was then investigated for leaf fungal communities. In the PCoAs, samples from conventional and organic sites did cluster independently across all sampling time-points (**Figure 3**). When all leaf samples were analyzed together, agricultural management strategy explained 13% of the variation among samples (**Table 2A**). When the three time-points were evaluated independently, a greater proportion of the variation was explained by the agricultural management strategy in July (32%) and August (27%) than in May (13%) (**Table 2B-D**). Site identity explained from 15 to 31% of the variation across sampling time-points (**Table 2**). With samples from Honeycrisp and Spartan only, fungal communities from different agricultural management strategy also clustered separately (**Figure S3**), as agricultural management strategy explained 14% of the variation (**Table S7**).

**Figure 3.**
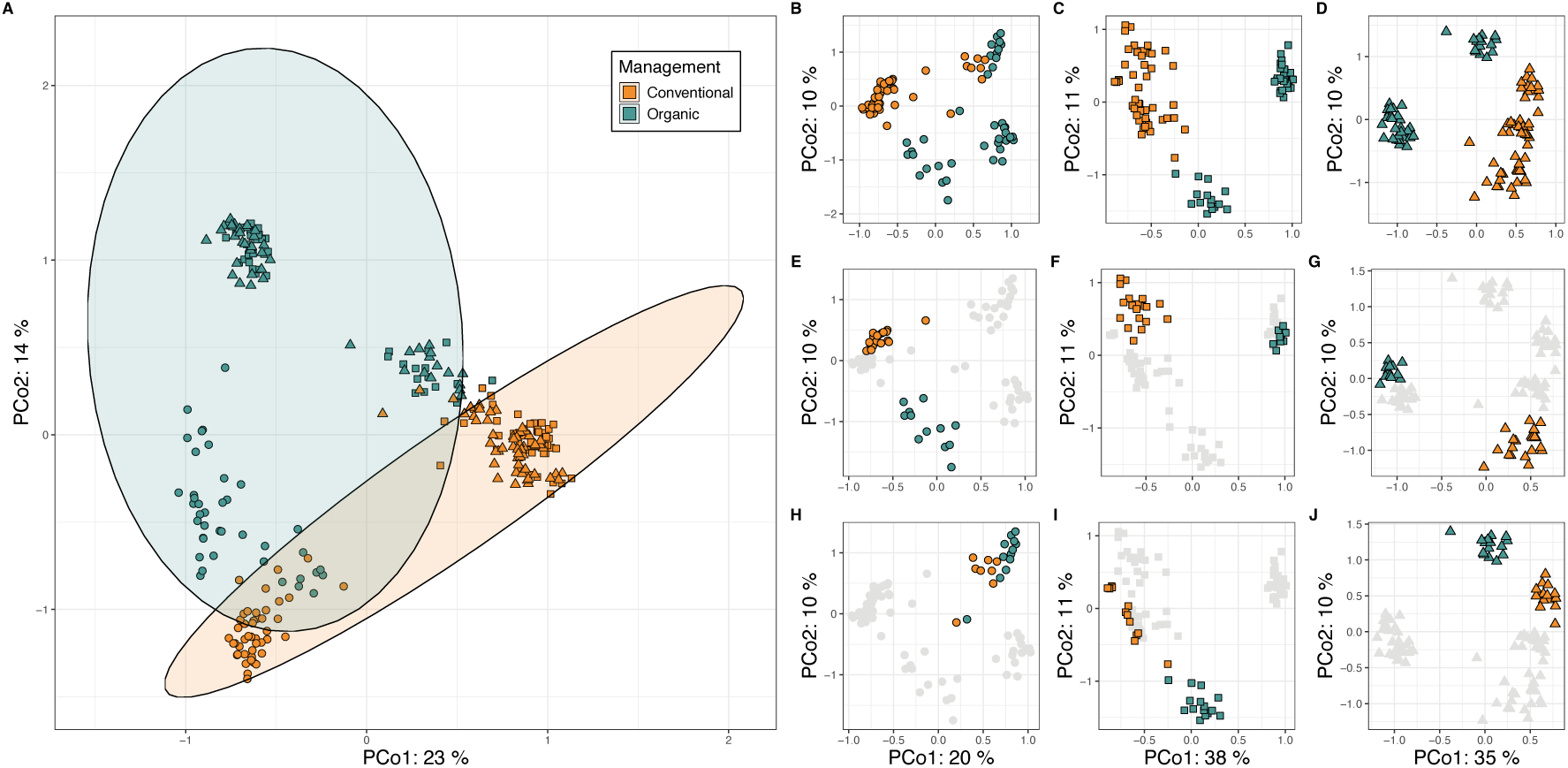
PCoAs on Bray-Curtis dissimilarities performed on leaf samples. **(A)** All leaf samples across sites and sampling months. **(B–D)** Subset of samples collected in May **(B)**, July **(C)**, and August **(D)**, with conventional sites shown in orange and organic sites in green. **(E–G)** Highlighted comparison of sites B1 (conventional) and B2 (organic for May **(E)**, July **(F)**, and August **(G)**; other sites are shown in gray. **(H–J)** Highlighted comparison of sites D1 (conventional) and D2 (organic) for May **(H)**, July **(I)**, and August **(J)**; other sites are shown in gray. Ellipses represent 95% confidence intervals.

**Table 2.** Factors explaining the variation within all leaf samples (PERMANOVA on Bray-Curtis dissimilarities). All leaf samples from all time-points (**A**) and separated for May (**B**), July (**C**) and August (**D**).

| **A** | R<sup>2</sup>(%) | F | Pr(>F) | **B** | R<sup>2</sup>(%) | F | Pr(>F) |
| --- | --- | --- | --- | --- | --- | --- | --- |
| Management strategy | 13.8 | 117.4 | 0.001 *** | Management strategy | 13.2 | 20.3 | 0.001 *** |
| Time | 19.1 | 81.1 | 0.001 *** | Cultivar | 10.2 | 2.2 | 0.001 *** |
| Cultivar | 4.4 | 5.3 | 0.001 *** | Management:Site*<sup>1</sup> | 24.6 | 9.5 | 0.001 *** |
| Management:Site*<sup>1</sup> | 15.4 | 32.7 | 0.001 *** | Management:Cultivar | 5.6 | 1.4 | 0.011 * |
| Management:Time | 6.0 | 25.7 | 0.001 *** | Management:Site:Cultivar | 12.2 | 1.4 | 0.001 *** |
| Management:Cultivar | 2.1 | 2.9 | 0.001 *** | Residuals | 34.3 |  |  |
| Management:Site:Time | 7.6 | 8.0 | 0.001 *** | Total explained | 65.7 |  |  |
| Management:Site:Cultivar | 4.5 | 2.8 | 0.001 *** |  |  |  |  |
| Residuals | 27.1 |  |  |  |  |  |  |
| Total explained | 72.9 |  |  |  |  |  |  |

| **C** | R<sup>2</sup>(%) | F | Pr(>F) | **D** | R<sup>2</sup>(%) | F | Pr(>F) |
| --- | --- | --- | --- | --- | --- | --- | --- |
| Management strategy | 32.6 | 108.4 | 0.001 *** | Management strategy | 27.5 | 88.5 | 0.001 *** |
| Cultivar | 8.3 | 3.9 | 0.001 *** | Cultivar | 8.5 | 3.9 | 0.001 *** |
| Management:Site*<sup>1</sup> | 26.6 | 22.1 | 0.001 *** | Management:Site*<sup>1</sup> | 31.1 | 25.0 | 0.001 *** |
| Management:Cultivar | 4.3 | 2.4 | 0.001 *** | Management:Cultivar | 4.1 | 2.2 | 0.003 ** |
| Management:Site:Cultivar | 10.4 | 2.5 | 0.001 *** | Management:Site:Cultivar | 8.6 | 2.0 | 0.001 *** |
| Residuals | 17.8 |  |  | Residuals | 20.2 |  |  |
| Total explained | 82.2 |  |  | Total explained | 79.8 |  |  |

\*<sup>1</sup>Site is nested within Management.

For leaves, fungal community composition was also explored across sampling time-points (May, July, and August) for both conventional and organic sites. The most relatively abundant fungal genera at conventional sites were *Sporobolomyces* (28%), *Aureobasidium* (15%), and *Cladosporium* (12%), summing up to 55% of sequences (**Figure 4**). Of note, the genus *Sporobolomyces* showed a strong increase in relative abundance at conventional sites from May (<1%) to July (51%) and August (32%) (**Figure 4**). The most relatively abundant fungal genera at organic sites were *Aureobasidium* (18%), *Didymella* (13%), and *Cladosporium* (11%), summing up to 42% of sequences (**Figure 4**). Similar community profiles were observed when analyzing the data subset including only Honeycrisp and Spartan cultivars (**Figure S4**).

**Figure 4.**
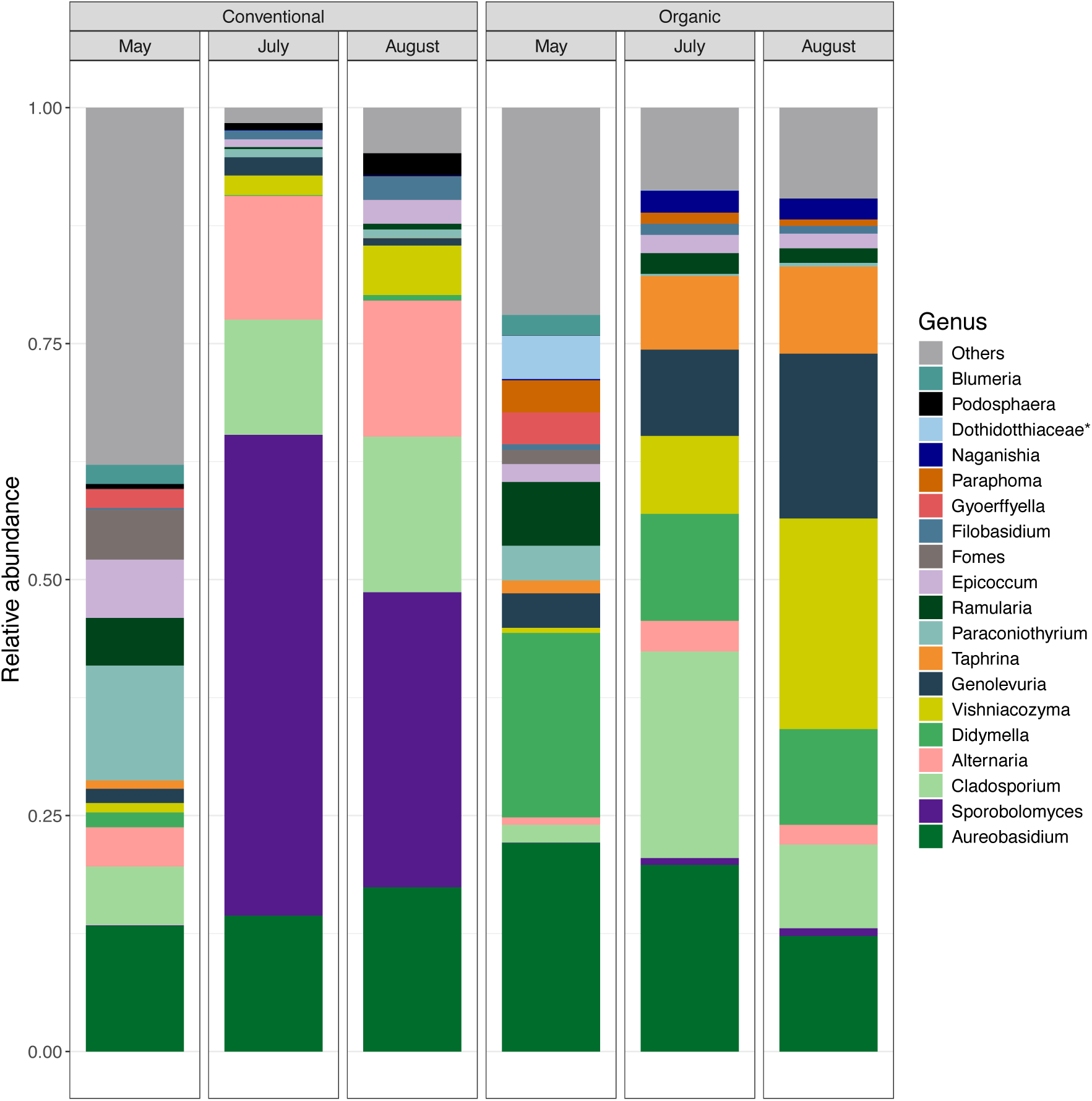
Fungal community composition across all leaf samples (all cultivars). Average relative abundance of the 20 most abundant fungal genera. Taxa not among the top 20 are grouped under “Others.” Bar charts are organized by agricultural management strategy and sampling month (Conventional: n_May_ = 45, n_July_ = 48, n_August_ = 54; Organic: n_May_ = 40, n_July_ = 44, n_August_ = 44). Asterisks indicate taxa identified at the family level but with unknown genus.

For both management strategies, our results show that there is a shift in fungal community composition between May and July (**Figure 4**), which is confirmed by the change in the absolute abundance of certain taxa (**Figure 5**) as well as by general changes in the relative abundance of specific fungal classes (**Figure S6**). In leaf samples, 10 and 28 genera were found to be differentially enriched in conventional and organic sites, respectively (**Figure 5**). Several fungal classes showed taxonomic coherence in their associations with conventional or organic management strategies. Notably, Lecanoromycetes genera *(Scoliciospurum* and *Hypoguymnia*), known to be lichen-forming or lichenicolous fungi, were identified as significantly enriched only with conventional sites (**Figure 5**). On the other hand, two classes of basidiomycetous yeasts, the Tremellomycetes (*Papiliotrema*, *Genolevuria*, *Dioszegia*, *Naganishia*, and *Vishniacozyma*) and Cystobasidiomycetes (*Cystobasidium*, *Symmetrospora*, and *Buckleyzyma*) were only enriched at organic sites (**Figure 5**). Of note, the potentially pathogenic genera *Podosphaera* and *Alternaria* were enriched at conventional orchards, while *Ramularia*, *Taphrina*, and *Didymella* were enriched at organic orchards (**Figure 5**). The most differentially enriched genus was an unnamed Dothidotthiaceae, estimated to be 75 times more abundant (4.32 ± 0.13 LFD) in organic sites compared to conventional sites. In terms of alpha diversity, a general decrease in Shannon index through time was observed at conventional sites (**Figure 6A-B**). Oppositely, alpha diversity remains stable from May to August at organic sites (**Figure 6A-B**). When comparing both management strategies through sampling month, alpha diversity was higher at conventional sites in May, while the opposite was observed in July and August with higher diversity at organic sites (**Figure 6C**).

**Figure 5.**
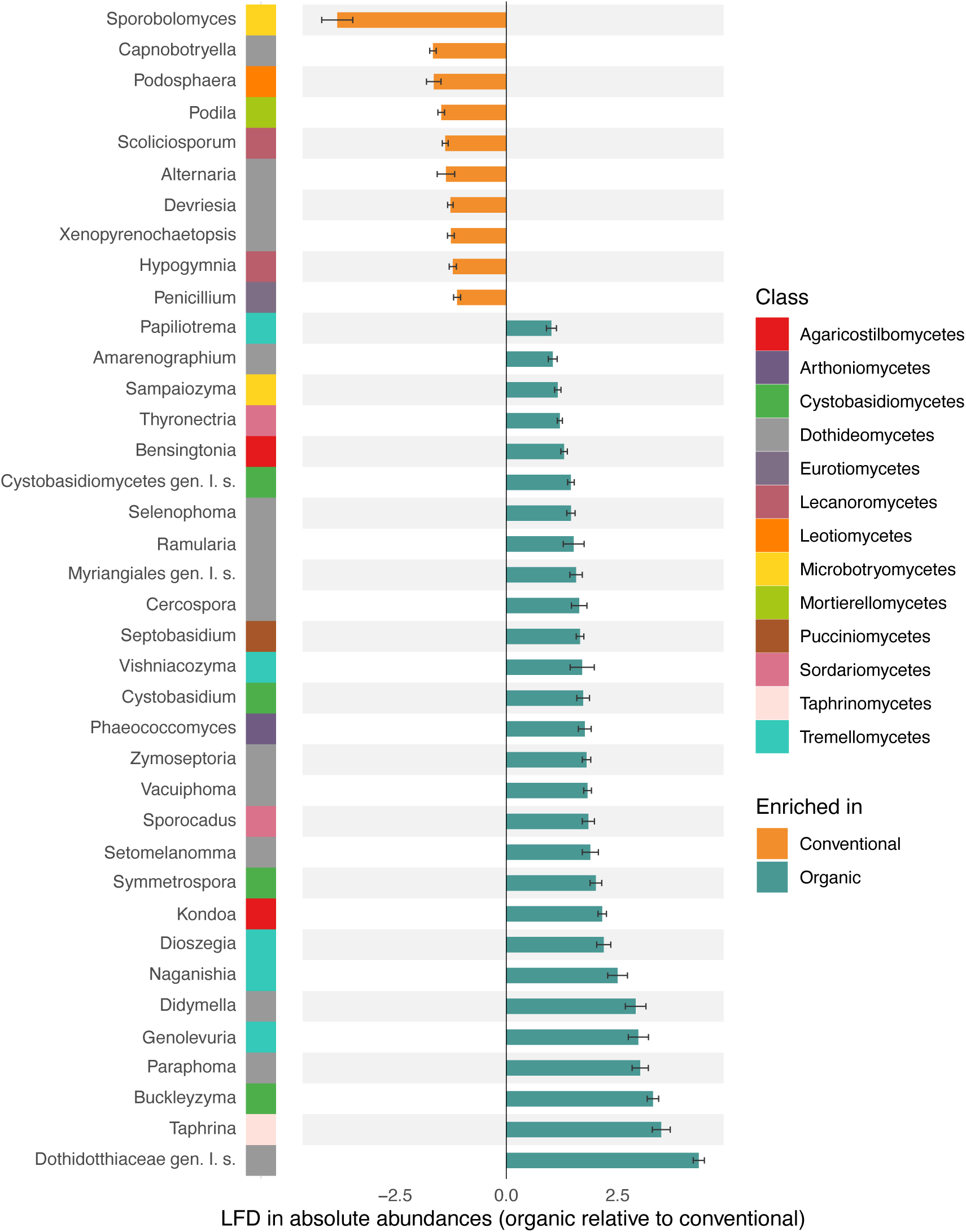
Differential abundance analysis of leaf communities comparing differences in absolute abundance of genera across agricultural management strategies, controlling for cultivar. A negative natural log fold difference (LFD) corresponds to an enrichment in the conventional sites (i.e., the reference group in the log-linear model), whereas a positive LFD corresponds to an enrichment in the organic group. Gen. I. s.: *Genus Incertae sedis*.

**Figure 6.**
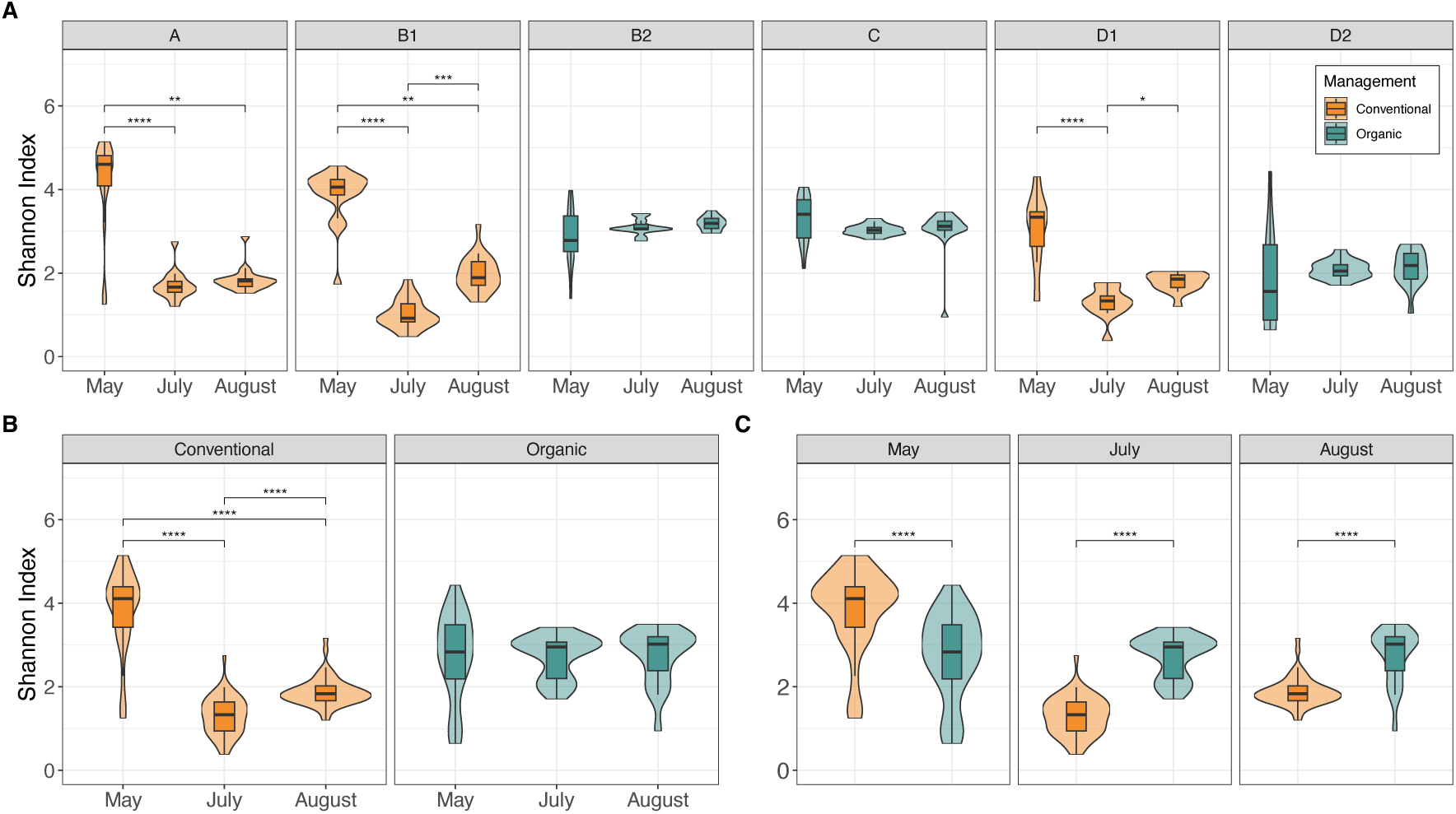
Shannon alpha diversity of leaf samples (all cultivars). **(A)** Alpha diversity across sampling times for each individual site. **(B)** Alpha diversity across sampling times grouped by agricultural management strategy. **(C)** Comparison of alpha diversity between conventional and organic management strategies at each sampling time point. Asterisks indicate significance levels from post-hoc Dunn tests following a Kruskal-Wallis test (*P* ≤ 0.05; **P** ≤ 0.01; ***P*** ≤ 0.001; **P** ≤ 0.0001).

## Discussion

Understanding the dynamics of microbial communities in agroecosystems is essential to support the improvement of management strategies towards more resilient and sustainable crop production. While many observational studies have focused on spatiotemporal variation in microbial communities [49, 50], few have examined these dynamics under *in field* conditions with direct relevance to producers. In this study, we extended the design of our previous works on the apple phyllosphere to assess how fungal communities respond to agricultural management strategy (conventional vs. organic) across multiple sites (six orchards) and time points (three sampling months).

Overall, our results broadly support our initial hypotheses. First, consistent with our expectation that fungal community composition would diverge between management strategies, our data shows that management strategy is associated with alterations in the composition and diversity of fungal communities of the apple tree flowers and leaves, with effects intensifying from May to July before stabilizing in August (**Tables 2-3**; **Figures 1-3**). The differential abundance of specific taxa between conventional and organic sites, including apple pathogens such as *V. inaequalis*, further supports the idea that selective pressure from agricultural chemical use (natural vs. synthetic) drives strong variations in dominant fungal taxa. Second, contrary to our expectation that alpha diversity would be reduced at conventional sites, our data showed time-dependent differences between management strategies, with higher diversity at conventional sites early in the season (May) but higher diversity at organic sites later in the season (July-August), alongside a general decline through time at conventional sites and a relative stability at organic sites. This pattern suggests that synthetic chemical inputs may initially allow for higher diversity but subsequently lead to a decline over the growing season, whereas organic management may promote more stable fungal alpha diversity dynamics throughout the growth season. Yet, more information about the exact products used and timing of applications is needed to link directly interventions to mycobiome dynamics. Third, and consistent with our expectation that spatial and temporal variation would be detected, both site and month of sampling were strongly correlated with changes in leaf fungal community composition (**Table 1**; **Table 2A**), reinforcing the idea that orchard-specific conditions, such as microclimate, local vegetation, and management strategies, contribute to shaping the local phyllosphere mycobiome, alongside temporal dynamics likely driven by climatic cues and dispersal from local reservoirs [51].

Even though agricultural management strategies were broadly categorized as conventional or organic due to limited access to detailed treatment records, this binary classification revealed clear and consistent differences in fungal communities between orchard types. Working *in field* inevitably introduces constraints, such as incomplete knowledge of specific inputs or treatment schedules, yet it also offers ecological relevance and a more direct applicability of findings for growers. Our results showed distinct fungal community profiles between conventional and organic orchards (**Figures 1-4**), suggesting that the use of synthetic versus organic inputs can exert differential selective pressures on the phyllosphere mycobiome, for example resulting in shifts in community composition. A potential explanation for the lower fungal diversity in flowers and leaves in May at organic sites lies in the contrasting disturbance regimes imposed by management strategies.

Conventional orchards often rely on synthetic fungicides with broad-spectrum activity, which could have initially suppressed dominant pathogenic taxa and reduced competitive exclusion, thereby creating a more even community structure and increasing local diversity at the beginning of the season [49]. In contrast, organic management typically employs the repeated use of non-specific contact treatments (e.g., sulfur, copper, biological controls) that may limit disease symptoms without fully eradicating certain fungal pathogens or epiphytes [50]. This partial control could allow a few well-adapted taxa to rapidly dominate the phyllosphere in May, reducing the overall diversity [51]. Early in the summer, organic management strategies may favor taxa with strong competitive abilities under lower chemical disturbance, allowing for a stronger role for ecological drivers such as priority effects and niche monopolization. Interestingly, this trend was reversed in July and August, as organic sites showed higher alpha diversity than conventional sites (**Figure 6**). Overall, our results suggest that management strategies and chemical selectivity differentially affect fungal communities through contrasting mechanisms of disturbance and competition. Although orchards were categorized as conventional or organic, management strategies in reality exist along a continuum, with many conventional producers integrating elements of organic strategies and carefully considering the ecological and health implications of synthetic inputs. This highlights the need for research on the development of adaptive, hybrid approaches that support both disease control and sustainable phyllosphere microbiome dynamics.

### Flower mycobiome of organic and conventional orchards host different plant pathogens

In flower samples, fungal communities from conventional and organic orchards formed distinct clusters in PCoA ordinations (**Figure 1A**). This pattern was supported by PERMANOVAs, indicating that agricultural management strategy explains a significant proportion of variation in flower fungal community composition. This observation is consistent with management-driven environmental filtering, whereby contrasting input regimes lead to different selection pressures on fungal taxa. In conventional systems, synthetic fungicide applications likely act as strong filters favoring tolerant or resistant taxa, whereas organic systems may maintain a pool of dominant taxa under comparatively weaker chemical filtering. For instance, genera such as *Didymella* and *Ramularia*, which include potential plant pathogens species (e.g., *R. collo-cygni*, *D. pomorum*) [52, 53], were more relatively abundant and differentially enriched on flowers at organic orchards (**Figures 1-2**). This pattern may reflect reduced suppression under lower chemical input regimes, allowing opportunistic or latent pathogens to persist within the community, as well as contributing to the lower diversity observed in organic flower samples (**Figure 1C**). In contrast, *Alternaria* and *Podosphaera*, which include potential plant pathogens species (e.g., *A. mali* and *P. leucotricha*) [24, 25] were more relatively abundant and differentially enriched at conventional sites compared to organic ones (**Figure 1**), suggesting selection for taxa tolerant to fungicide exposure.

### The effect of management strategy on leaf mycobiome beta diversity increases with time

Similar selection patterns were observed in leaf samples. Management strategy associated with shifts in fungal composition and diversity, which intensified over the growing season. The increasing proportion of variance explained by management (from 13% in May to 32% in July and 27% in August; **Table 2**) suggests a temporal strengthening of environmental filtering, potentially driven by the cumulative effects of repeated treatments. Early-season communities may be largely shaped by dispersal from overwintering reservoirs and initial colonization (i.e., priority effects), whereas later-season communities could increasingly reflect selection and microbe-microbe interactions under contrasting management regimes. In conventional orchards, the mid-season increase in *Sporobolomyces*’ relative abundance (**Figure 4**), may indicate either higher tolerance to treatment or a release from competition under selective pressure. Given that members of this genus produce carotenoids with antimicrobial properties (e.g., red pigmented *Sporobolomyces*) and thus have a potential for biocontrol [54, 55], it is commonly associated with beneficial or commensal roles in the phyllosphere [56–58]. Their proliferation may further influence community assembly through interference competition, contributing to the observed decline in alpha diversity later in the season (**Figure 6**). On the other hand, the dominance of *Cladosporium* and *Aureobasidium* across both management strategies (**Figure 4**) suggests that these genera are either highly resilient to environmental filtering or maintained through repeated dispersal and application. Different species of *Cladosporium* have shown biocontrol potential against plant pathogens [56, 59]. In particular, the widespread use of *Aureobasidium*-based biocontrol products (e.g., Blossom Protect) likely reinforces its persistence across sites, illustrating how management practices can directly shape leaf community composition [9]. Finally, the higher abundance of Dothideomycetes at organic sites may reflect weaker suppression of latent pathogens and early colonizing saprotrophs [60], reinforcing the idea that management influences not only diversity but also the balance among functional groups within the phyllosphere. These shifts in dominant taxa likely influence community structure through competitive interactions, with the eventual dominance by a few taxa potentially leading to a lower alpha diversity at conventional sites [61, 62]. At organic sites, the coexistence of multiple taxa, including saprotrophs such as *Exidia* (Agaricomycetes, known as saprotroph jelly fungi), suggests weaker filtering. Importantly, several of the genera found here (e.g., *Alternaria*, *Didymella*, *Ramularia*, and *Taphrina*) are known to exhibit flexible colonization strategies, including epiphytic, endophytic, and pathogenic phases depending on environmental conditions and host status, reflecting the broader capacity of many fungi to shift along a symbiosis-pathogenicity continuum [63]. Further research, explicitly disentangling leaf epiphytes from endophytes, is needed to test if the shifts in leaf fungal community composition across management strategies also impact mycobiome functional diversity.

Overall, these findings highlight how overlapping and distinct management interventions across organic and conventional regimes impose unique selection pressures on phyllosphere fungal communities, leading to both shared dominant taxa, such as *Aureobasidium*, and divergent overall community structures. In terms of diversity, alpha diversity declined over time in conventional orchards but remained stable in organic sites, a pattern consistent with our previous observations [26]. Leaf fungal alpha diversity was higher at conventional sites in May, while organic sites showed greater diversity in July and August. It is interesting to note that, for bacteria, we had also observed a lower alpha diversity at organic sites in May compared to conventional sites [27]. Taken together, our results provide *in field* information about apple tree flower and leaf mycobiome dynamics highlighting that agricultural management strategies are associated with shifts in fungal community composition and diversity across the growing season. By linking fungal dynamics to orchard management, this work helps better understand how management strategies shape phyllosphere fungal communities, offering a foundation for future research aimed at developing mycobiome-informed strategies to support plant health and disease resistance. Further work is needed to increase spatial and temporal resolutions while incorporating more information about timing of plant protection product applications.

### Apple tree cultivar is weakly associated with fungal phyllosphere communities

Consistent with earlier observations by Boutin et al. (2024) [26], cultivar explained a small but significant portion of the variation in leaf-associated fungal community structure (**Table 2**). This modest effect aligns with previous studies showing that host cultivar identity tends to exert a smaller influence on microbial communities than host species identity [49, 64, 65]. Nonetheless, the reproducibility of this pattern across independent datasets underscores that, although this effect is weaker than month of sampling, site identity, or agricultural management strategies, accounting for cultivar identity is relevant to understand the dynamics of the apple phyllosphere mycobiome.

### Limitations

While our study provides ecologically relevant insights into the influence of management strategies on apple tree phyllosphere fungal communities, several limitations should be acknowledged. First, working under *in field* conditions meant that we could not fully control environmental variables or access detailed treatment records for each orchard (e.g., no access to flowers at sites D1 and D2, cultivars not accessible at all sites, approximation of BBCH exact stage at sampling). As a result, agricultural management strategies were broadly categorized as conventional or organic, which may mask important nuances in management strategies and input types. Organic orchards were managed under Canadian Ecocert certification standards, which restrict the use of synthetic fertilizers and pesticides while allowing a defined set of inputs such as biocontrol agents, mineral-based products (e.g., sulfur, copper), and certain organically approved compounds. Although this framework provides a consistent regulatory baseline, inputs and application frequencies can still vary among producers, potentially contributing to within-category heterogeneity.

Second, our sequencing approach targeted the ITS1 region, which, while widely used for fungal community profiling, has known biases in taxonomic resolution and amplification efficiency across fungal taxa. These methodological constraints may have influenced the detection and relative abundance of certain groups, which should be held into account when interpreting or comparing the results with other studies. In addition, DNA sequencing does not necessarily reflect gene activity nor microbial viability. Third, although this study focussed on the phyllosphere, future research integrating both rhizosphere and phyllosphere perspectives are needed, as their combined influence on plant fitness and resilience is critical to optimizing sustainable management strategies. Finally, although our study design captured temporal and spatial variation, it remains observational in nature and cannot establish causal relationships between specific treatments and microbial outcomes. Future studies, covering multiple years, incorporating controlled experiments, metagenomic approaches, and detailed agronomic metadata will be essential to refine our understanding of how orchard management shapes fungal community structure and function.

### Conclusion

This study provides novel information about how agricultural management strategies are associated with fungal community composition and diversity of the apple tree phyllosphere under *in field* conditions. By leveraging a robust sampling design across multiple sites, cultivars, and months of sampling, we were able to quantify the relative influence of agricultural management strategies, time, and site on fungal communities, track alpha diversity dynamics throughout the growing season, and identify differentially abundant taxa associated with each management strategy. Our findings offer new insights into how agricultural management strategies used for pathogen control may differentially affect the structure of phyllosphere fungal communities, including shifts in relative abundance of both pathogenic and beneficial taxa. Beyond documenting these patterns, our work contributes to a growing body of research emphasizing the ecological relevance of phyllosphere fungi, which play key roles in plant health, disease resistance, and microbial interactions. Studying these communities *in situ* highlights how site-level decisions can influence microbial composition and, ultimately, crop resilience and productivity.

While our results reveal clear differences between conventional and organic systems, determining which management strategy is more sustainable remains an open question. Future studies are required to develop integrative sustainability indices and link microbial community structure to disease incidence, yield, and broader environmental and socio-economic outcomes. We therefore advocate for future research that bridges microbial ecology with on-farms agronomic performance and sustainability science to guide the development of microbiome-informed orchard management strategies.

## Acknowledgments

We want to acknowledge the help of the Laforest-Lapointe lab members who participated to field work (E. Lussier, G. Lajeunesse, M. Enea, J. Fraysse, A. Werz, R. Perreault) and revised the manuscript (E. Lussier). We also want to thank orchards managers who allowed us to go sampling on their land.

## Authors contribution

Conceptualization: S.B. & I.L-L.

Data curation: S.B., J.R-L. & A.R.

Formal analysis: A.R., S.B, J.R-L & I.L-L.

Funding acquisition: I.L-L., S.B. & J.R-L.

Investigation: S.B., J.R-L. & A.R.

Methodology: S.B., J.R-L. & A.R.

Project administration: I.L-L.

Resources: I.L-L.

Software: S.B., J.R-L. & A.R.

Supervision: I.L-L.

Validation: J.R-L.

Visualization: S.B., J.R-L., A.R. & I.L-L.

Writing – original draft: S.B., J.R-L., A.R. & I.L-L.

Writing – review & editing: I.L-L. & J.R-L.

## Data availability

The data underlying this article are available in European Nucleotide Archive at https://www.ebi.ac.uk/ena, and can be accessed with PRJEB96463. All scripts are available at the following GitHub repository: https://github.com/alexis-roy5/orchard_phyllosphere

