## Supplementary tables and figures for "A Field-Based Study of Phyllosphere Mycobiomes in Apple Orchards Under Varying Agricultural Management Strategies"

**Table S1.** Studies investigating the impact of agricultural management strategies on phyllosphere bacterial and fungal communities.

| Reference | Plant | Type of study | Impact of agricultural strategies on diversity | | | |
|---|---|---|---|---|---|---|
| | | | Bacteria | | Fungi | |
| | | | α | β | α | β |
| Karlsson et al. (2017) | Wheat | Field | NA | NA | Yes* | Yes |
| Castañeda et al. (2018) | Grapevine | Field | NA | NA | No | Yes |
| Knorr et al. (2019) | Wheat | Field | NA | NA | No | Yes |
| Katsoula et al. (2020) | Pepper | Field | No | Yes | Yes | Yes |
| Wu et al. (2023) | Cucumber | Greenhouse | No | No | No | Yes |

*Richness

**Table S2.** Description of cultivars sampled at each site.

| SITES | Strategies | CULTIVARS | | | | | | | |
|---|---|---|---|---|---|---|---|---|---|
| | | Cortland | Paulared | Honeycrisp | Spartan | Sunrise | Liberty | Empire | Freedom |
| A | Conventional | X | X | X | X | | X | X | |
| B1 | Conventional | X | X | X | X | X | X | X | |
| B2 | Organic | | | X | X | X | | X | |
| C | Organic | X | X | X | X | | X | | X |
| D1 | Conventional | X | | X | X | | X | X | |
| D2 | Organic | X | | X | X | | X | X | |

**Table S3.** Metadata of samples removed during bioinformatic processing.

| Sample | Time | Code | Strategy | Cultivar | Type | Tree replicate | Sequences |
|---|---|---|---|---|---|---|---|
| 2023-1-ASB-CL-LE-1 | May | A | Conventional | Cortland | Leaf | 1 | 41 |
| 2023-1-ASB-EM-FL-2 | May | A | Conventional | Empire | Flower | 2 | 518 |
| 2023-1-ASB-HC-FL-1 | May | A | Conventional | Honeycrisp | Flower | 1 | 226 |
| 2023-1-ASB-LI-FL-3 | May | A | Conventional | Liberty | Flower | 3 | 1347 |
| 2023-1-COM-EM-FL-3 | May | B2 | Organic | Empire | Flower | 3 | 0 |
| 2023-1-COM-SP-FL-1 | May | B2 | Organic | Spartan | Flower | 1 | 2 |
| 2023-1-COM-SP-FL-3 | May | B2 | Organic | Spartan | Flower | 3 | 38 |
| 2023-1-COM-SR-FL-1 | May | B2 | Organic | Sunrise | Flower | 1 | 3 |
| 2023-1-COM-SR-FL-3 | May | B2 | Organic | Sunrise | Flower | 3 | 1255 |
| 2023-1-MIB-CL-LE-1 | May | D2 | Organic | Cortland | Leaf | 1 | 679 |
| 2023-1-MIB-EM-LE-3 | May | D2 | Organic | Empire | Leaf | 3 | 881 |
| 2023-1-MIB-LI-LE-1 | May | D2 | Organic | Liberty | Leaf | 1 | 14 |
| 2023-1-MIC-CL-LE-1 | May | D1 | Conventional | Cortland | Leaf | 1 | 390 |
| 2023-1-MIC-CL-LE-2 | May | D1 | Conventional | Cortland | Leaf | 2 | 1050 |
| 2023-1-MIC-CL-LE-3 | May | D1 | Conventional | Cortland | Leaf | 3 | 4 |
| 2023-1-MIC-EM-LE-1 | May | D1 | Conventional | Empire | Leaf | 1 | 6 |
| 2023-1-MIC-EM-LE-3 | May | D1 | Conventional | Empire | Leaf | 3 | 26 |
| 2023-1-MIC-LI-LE-2 | May | D1 | Conventional | Liberty | Leaf | 2 | 690 |
| 2023-1-MIC-LI-LE-3 | May | D1 | Conventional | Liberty | Leaf | 3 | 1404 |
| 2023-1-PMB-CL-FL-2 | May | B1 | Conventional | Cortland | Flower | 2 | 0 |
| 2023-1-PMB-LI-LE-2 | May | B1 | Conventional | Liberty | Leaf | 2 | 179 |
| 2023-1-VBS-LI-FL-2 | May | C | Organic | Liberty | Flower | 2 | 3 |
| 2023-1-VBS-LI-FL-3 | May | C | Organic | Liberty | Flower | 3 | 5 |
| 2023-1-VBS-LI-LE-1 | May | C | Organic | Liberty | Leaf | 1 | 1 |
| 2023-1-VBS-SP-FL-3 | May | C | Organic | Spartan | Flower | 3 | 2039 |
| 2023-1-VBS-SP-LE-2 | May | C | Organic | Spartan | Leaf | 2 | 8 |
| 2023-2-ASB-CL-LE-3 | July | A | Conventional | Cortland | Leaf | 3 | 16 |
| 2023-2-ASB-HC-LE-2 | July | A | Conventional | Honeycrisp | Leaf | 2 | 97 |
| 2023-2-ASB-PR-LE-3 | July | A | Conventional | Paulared | Leaf | 3 | 1563 |
| 2023-2-MIC-CL-LE-1 | July | D1 | Conventional | Cortland | Leaf | 1 | 30 |
| 2023-2-MIC-CL-LE-3 | July | D1 | Conventional | Cortland | Leaf | 3 | 39 |
| 2023-2-MIC-HC-LE-2 | July | D1 | Conventional | Honeycrisp | Leaf | 2 | 25 |
| 2023-2-VBS-CL-LE-3 | July | C | Organic | Cortland | Leaf | 3 | 6 |
| 2023-3-VBS-HC-LE-3 | August | C | Organic | Honeycrisp | Leaf | 3 | 1 |

**Table S4.** Pre- and post-rarefaction dataset descriptive statistics.

| | | ITS | ITS_rarefied |
|---|---|---|---|
| Total # of | Sequence | 5,022,964 | 822,500 |
| | ASV | 2,812 | 2,795 |
| | Sample | 329 | 329 |
| Sequence per sample | Mean | 15,267 | 2,500 |
| | SD | 8,189 | 0 |
| | Min | 2,508 | 2,500 |
| | Max | 47,875 | 2,500 |
| ASV per sample | Mean | 216 | 146 |
| | SD | 144 | 112 |
| | Min | 22 | 12 |
| | Max | 643 | 437 |
| ASV prevalence | Mean | 25 | 17 |
| | SD | 39 | 31 |
| | Min | 1 | 1 |
| | Max | 329 | 329 |

**Table S5.** Results of the tests for homogeneity of variance (*betadisper*) across groups for each PERMANOVA variables and their corresponding tables.

| Table | Data subset | Variable | Groups | F | P | Homogeneity |
|---|---|---|---|---|---|---|
| 1 | Flowers (all cultivars) | Strategy | 2 | 10.091 | 0.0025 | No |
| | | Cultivar | 8 | 11.690 | <0.0001 | No |
| | | Site | 4 | 34.338 | <0.0001 | No |
| S1 | Flowers (Honeycrisp & Spartan) | Strategy | 2 | 54.635 | 0.0000 | No |
| | | Cultivar | 2 | 0.001 | 0.9825 | Yes |
| | | Site | 3 | 39.738 | <0.0001 | No |
| 2A | Leaves (all cultivars) | Strategy | 2 | 0.002 | 0.9646 | Yes |
| | | Time | 3 | 5.769 | 0.0035 | No |
| | | Cultivar | 8 | 4.114 | 0.0003 | No |
| | | Site | 6 | 3.274 | 0.0069 | No |
| 2B | Leaves (May) | Strategy | 2 | 3.324 | 0.0719 | Yes |
| | | Cultivar | 8 | 5.497 | <0.0001 | No |
| | | Site | 6 | 2.488 | 0.0381 | No |
| 2C | Leaves (July) | Strategy | 2 | 12.768 | 0.0006 | No |
| | | Cultivar | 8 | 7.321 | <0.0001 | No |
| | | Site | 6 | 3.133 | 0.0120 | Yes |
| 2D | Leaves (August) | Strategy | 2 | 26.373 | <0.0001 | No |
| | | Cultivar | 8 | 3.588 | 0.0019 | No |
| | | Site | 6 | 4.787 | 0.0006 | No |
| S2 | Leaves (Honeycrisp & Spartan) | Strategy | 2 | 0.857 | 0.3569 | Yes |
| | | Time | 3 | 2.154 | 0.1213 | Yes |
| | | Cultivar | 2 | 0.090 | 0.7649 | Yes |
| | | Site | 6 | 2.551 | 0.0325 | No |

**Table S6.** Factors explaining the variation within Honeycrisp and Spartan flower samples (PERMANOVA on Bray-Curtis dissimilarities)

| | $R^2$(%) | F | Pr(>F) | |
|---|---|---|---|---|
| Strategy | 29.2 | 6.8 | 0.001 | *** |
| Cultivar | 5.3 | 1.2 | 0.235 | ns |
| Strategy:Site[*1] | 11.9 | 2.8 | 0.015 | * |
| Strategy:Cultivar | 4.9 | 1.1 | 0.252 | ns |
| Strategy:Site:Cultivar | 5.6 | 1.3 | 0.177 | ns |
| Residuals | 43.2 | | | |
| Total explained | 56.8 | | | |

[*1] Site is nested within Strategy

**Table S7.** Factors explaining the variation within Honeycrisp and Spartan leaf samples (PERMANOVA on Bray-Curtis dissimilarities)

| | $R^2$(%) | F | Pr(>F) | |
|---|---|---|---|---|
| Strategy | 14.1 | 49.9 | 0.001 | *** |
| Time | 20.6 | 36.5 | 0.001 | *** |
| Cultivar | 1.3 | 4.5 | 0.001 | *** |
| Strategy:Site[*1] | 20.2 | 17.9 | 0.001 | *** |
| Strategy:Time | 6.6 | 11.7 | 0.001 | *** |
| Strategy:Cultivar | 0.7 | 2.5 | 0.001 | *** |
| Strategy:Site:Time | 10.5 | 4.6 | 0.001 | *** |
| Strategy:Site:Cultivar | 3.4 | 3.0 | 0.001 | *** |
| Residuals | 22.6 | | | |
| Total explained | 77.4 | | | |

**SUPPLEMENTARY FIGURES**

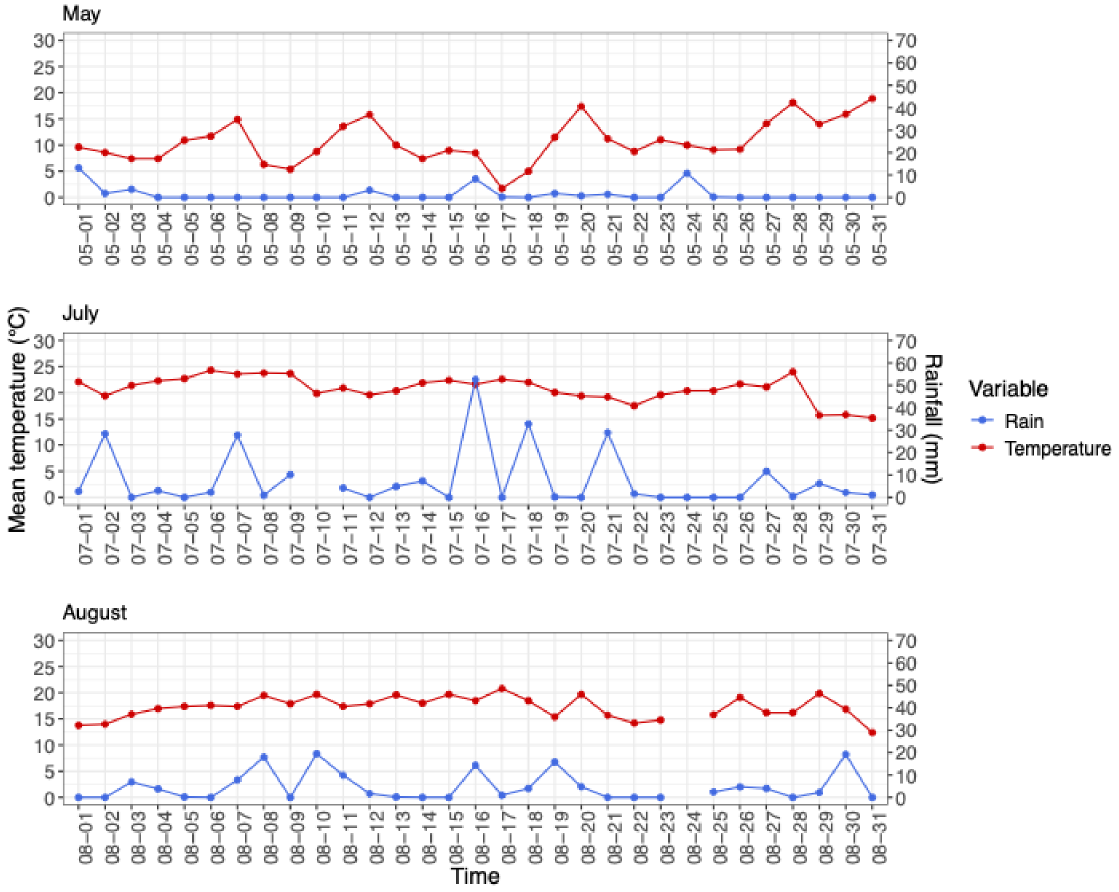


**Figure S1.** Mean temperature and total precipitation in May, July, and August 2023 at Sherbrooke meteorological station. Data retrieved from: https://climat.meteo.gc.ca/historical_data/search_historic_data_f.html

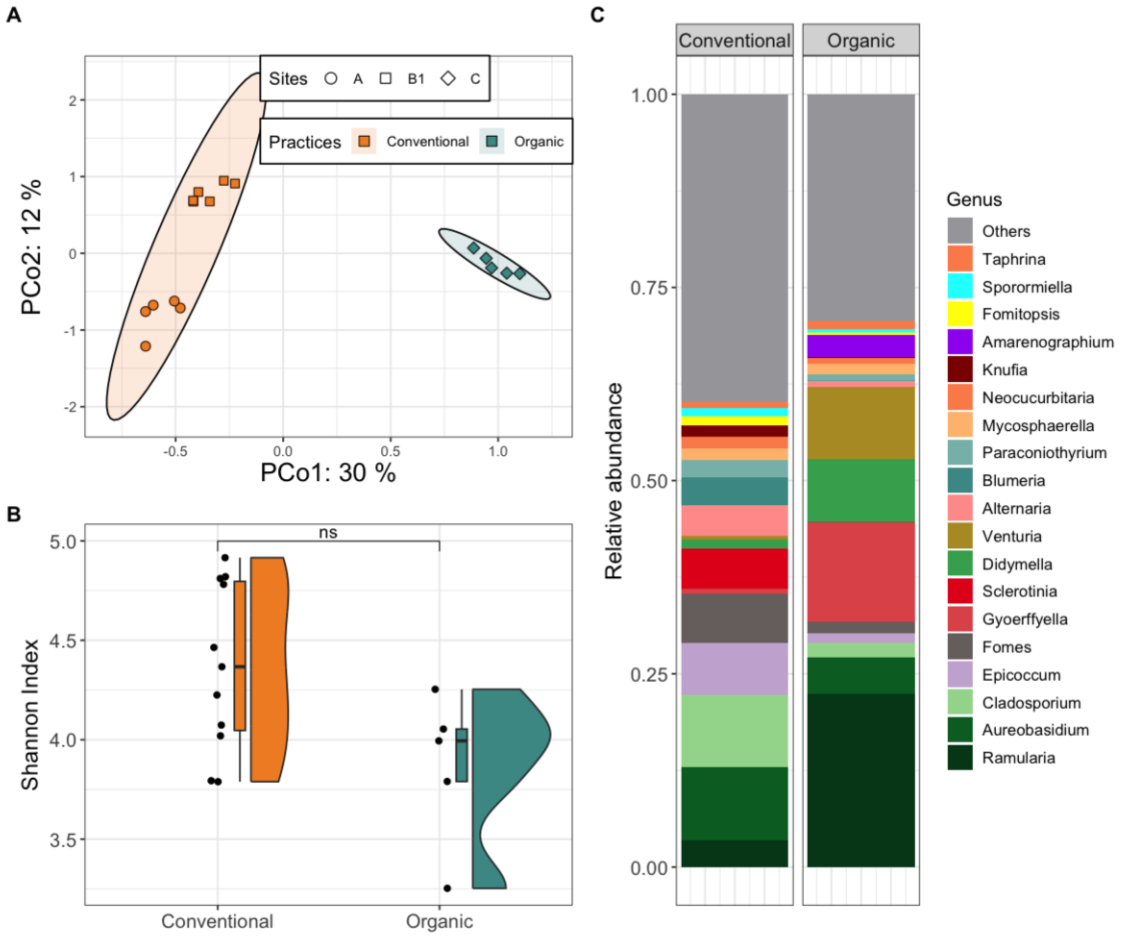


**Figure S2.** Flower Fungal composition and diversity from *Honeycrisp* and *Spartan* cultivars. **(A)** Principal Coordinates Analysis (PCoA) based on Bray-Curtis dissimilarities among flower samples. Point shapes and colors indicate site and agricultural Strategy. Ellipses represent 95% confidence intervals. **(B)** Shannon diversity index across conventional (n = 11) and organic (n = 5) sites. Asterisks denote significance levels from Wilcoxon-Mann-Whitney (*ns*, P > 0.05). **(C)** Average relative abundance of the 20 most abundant fungal genera across flower samples. Taxa not among the top 20 are grouped under "Others." Bar charts are separated by agricultural Strategy.

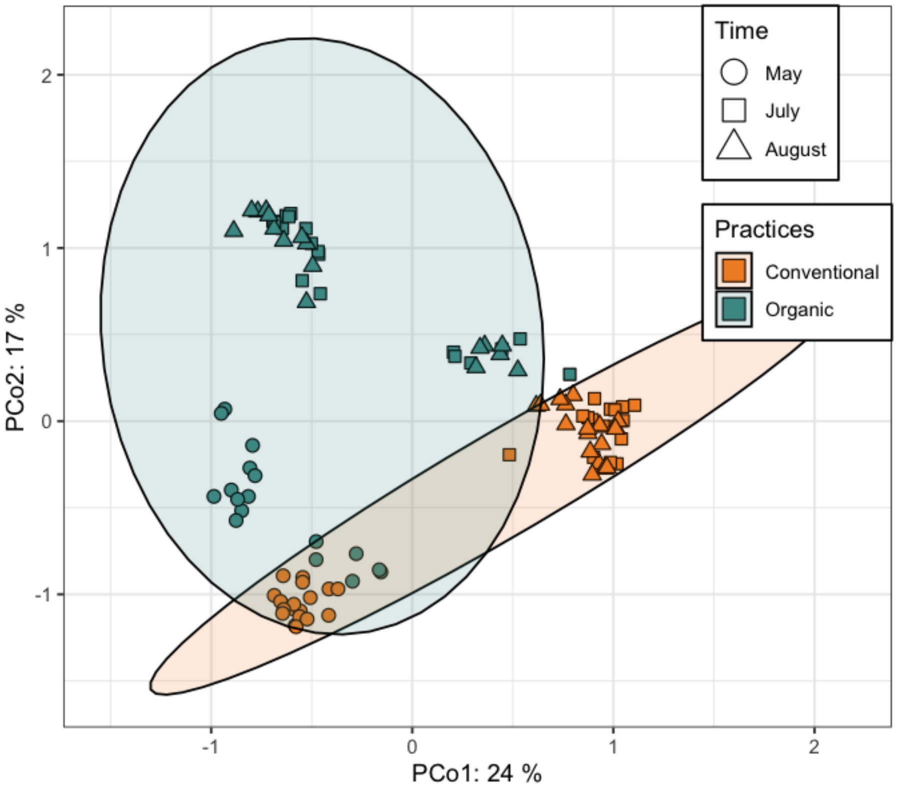


**Figure S3.** Principal Coordinates Analysis (PCoA) based on Bray-Curtis dissimilarities among leaf samples from *Honeycrisp* and *Spartan* cultivars. Point shapes represent sampling time, while point colors indicate agricultural management strategies. Ellipses denote 95% confidence intervals.

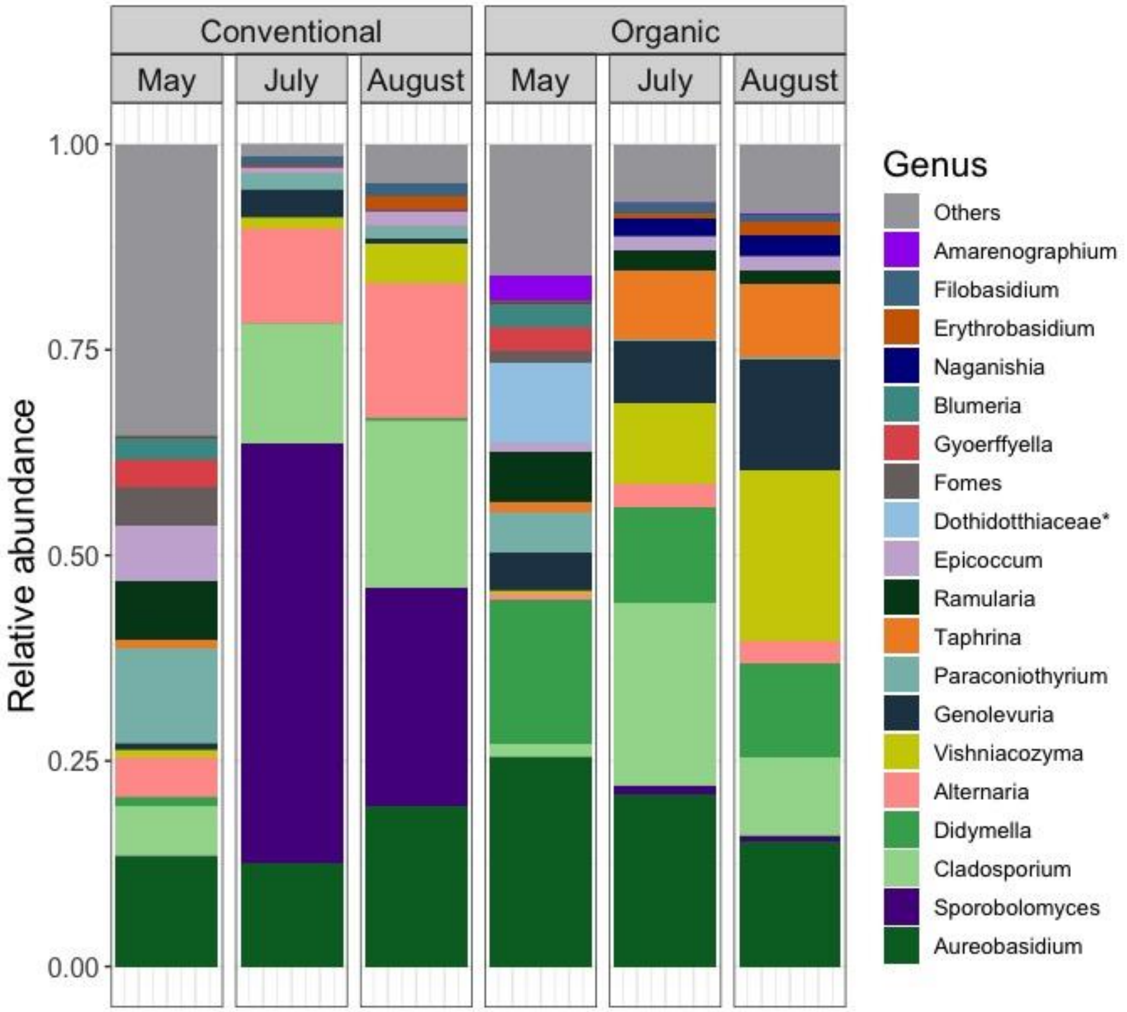


**Figure S4.** Fungal community composition in leaf samples from *Honeycrisp* and *Spartan* cultivars. Average relative abundance of the 20 most abundant fungal genera. Taxa not among the top 20 are grouped under "Others." Bar charts are organized by agricultural strategy and sampling month (Conventional: $n_{May}$ = 18, $n_{July}$ = 16, $n_{August}$ = 18; Organic: $n_{May}$ = 17, $n_{July}$ = 18, $n_{August}$ = 17). Asterisks indicate taxa identified at the family level but with unknown genus.

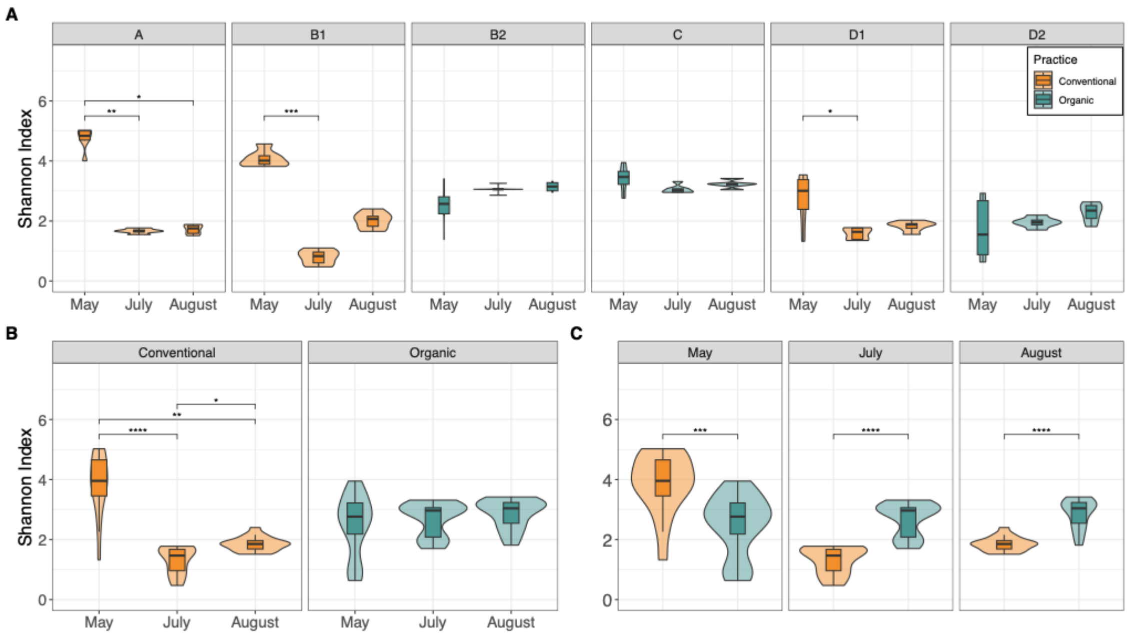


**Figure S5.** Shannon alpha diversity of leaf samples from *Honeycrisp* and *Spartan* cultivars. **(A)** Alpha diversity across sampling times for each individual site. **(B)** Alpha diversity across sampling times grouped by agricultural strategy. **(C)** Comparison of alpha diversity between conventional and organic strategies at each sampling time point. Asterisks indicate significance levels from post-hoc Dunn tests following a Kruskal-Wallis test ($P \leq 0.05$; **P** ≤ 0.01; ***P*** ≤ 0.001; **P** ≤ 0.0001).

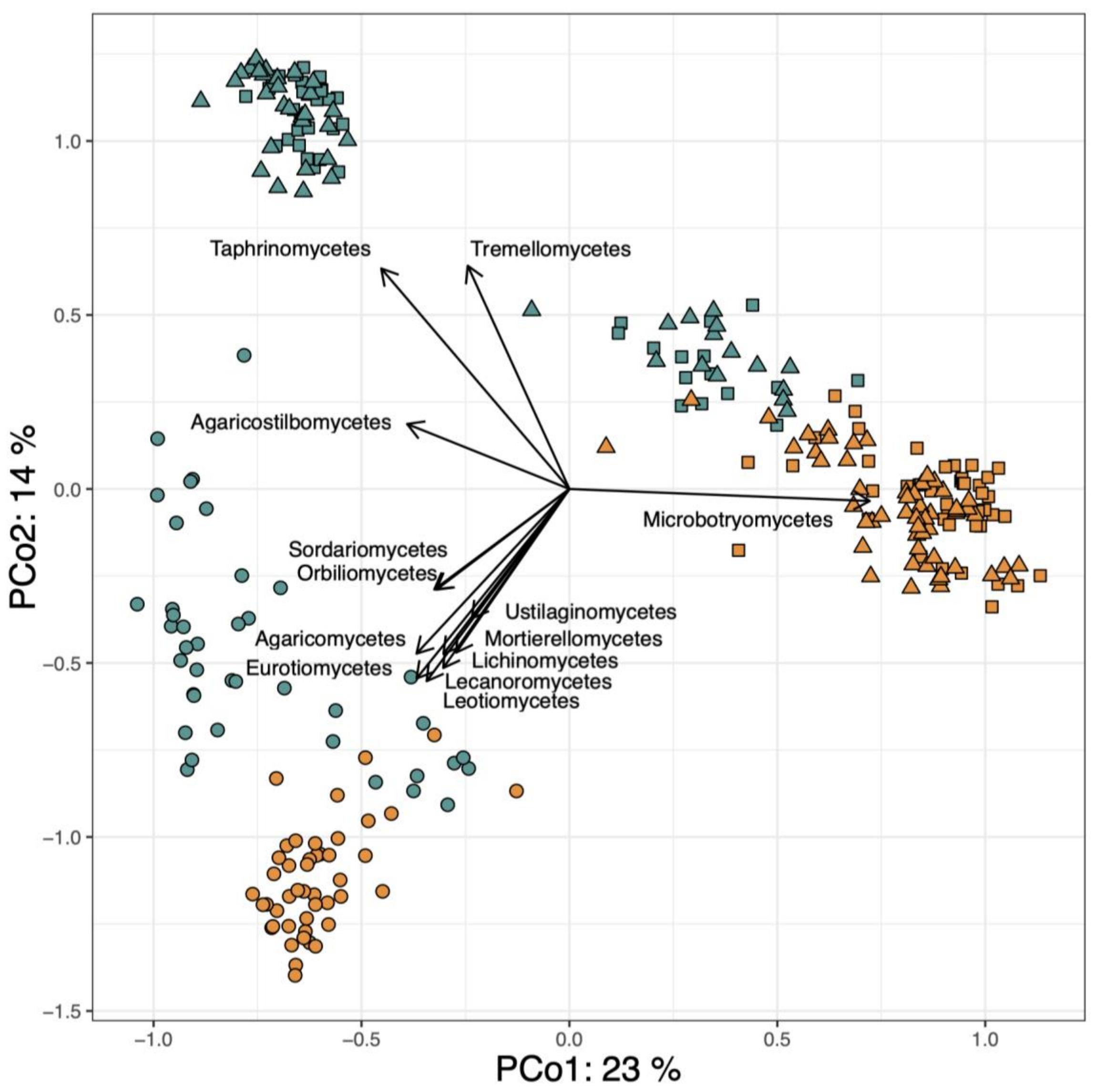


**Figure S6.** Principal Coordinates Analysis (PCoA) based on Bray-Curtis dissimilarities among phyllosphere samples collected from apple trees under conventional and organic agricultural management strategies. Each point represents a sample, colored by agricultural strategy. Arrows along the plot margins indicate fungal classes whose relative abundances are significantly correlated ($p < 0.001$, Bonferroni-corrected) with the ordination axes, as determined by the function *envfit*.
